# The effects of social experience on host gut microbiome in male zebrafish (*Danio rerio*)

**DOI:** 10.1101/2023.01.23.525298

**Authors:** Emily Scott, Michael S Brewer, Ariane L Peralta, Fadi A Issa

## Abstract

Although the gut and the brain vastly differ in physiological function, they have been interlinked in a variety of different neurological and behavioral disorders. The bacteria that comprise the gut microbiome communicate and influence the function of various physiological processes within the body including nervous system function. However, the effects of social experience in the context of dominance and chronic stress on gut microbiome remain poorly understood. Here, we examined whether social experience impacts the host zebrafish (*Danio rerio*) gut microbiome. We studied how social dominance during the first two weeks of social interactions changed the composition of zebrafish gut microbiome by comparing gut bacterial composition, diversity and relative abundance among socially dominant, submissive, social isolates, and control group-housed communal fish. Using amplicon sequencing of the 16S rRNA gene, we report that social dominance significantly affects host gut bacterial community composition but not bacterial diversity. At the genus-level, *Aeromonas* and unclassified Enterobacteriaceae relative abundance decreased in dominant individuals while commensal bacteria (e.g., *Exiguobacterium* and *Cetobacterium)* increased in relative abundance. Conversely, the relative abundance of *Psychrobacter* and *Acinetobacter* was increased in subordinates, isolates, and communal fish compared to dominant fish. The shift in commensal and pathogenic bacteria highlights the impact of social experience and the accompanying stress on gut microbiome with potentially similar effects in other social organisms.

**IMPORTANCE:** Disruptions in the gut microbiome negatively impact various systems in the body. Recently, the gut microbiome has been associated with neurological deficits in both behavioral and neurodegenerative disorders. Given the increasing prevalence in diagnosis of both neurological disease and behavioral disorders, researching the link between social behaviors and the gut microbiome is critical to better understand how the gut and the brain communicate during healthy and diseased states. Our research findings demonstrate the effects of social dominance and chronic stress on host gut microbiome composition. By identifying variations in bacterial relative abundance based on social experience and associated stress, a broader understanding of *pathogenic* (e.g., Enterobacteriaceae, *Aeromonas*) versus commensal communities (e.g., *Exiguobacterium, Cetobacterium*) and related host physiology can be inferred.

## INTRODUCTION

The gut microbiome plays key roles in biochemical functions in vertebrates. Gut bacterial metabolism has been linked to many systemic diseases in humans and influences numerous immune system pathways (1–5). Additionally, commensal bacteria within the gut microbiome serves as a line of defense to viral infections through interferon signaling (6). Along with these known functions of the gut biome, it has recently been linked to affecting brain function in the context of neurodevelopmental (7, 8), behavioral (9–11), and neurodegenerative disorders (12– 14). For instance, the gut microbiome produces certain biochemicals (i.e., dopamine and oxytocin) that regulate neural nuclei involved in social behavior (15, 16). This link is known as the “microbiota-gut-brain axis” and its dysfunction has been associated with many behavioral disorders such as autism spectrum disorder, depression, and anxiety with long-lasting negative impact on health and wellbeing (17). Thus, understanding the mechanisms of disease onset and progression related to the gut microbiome is critical.

The typical gut microbiome consists of both commensal and pathogenic bacteria (18– 20). While it is known that pathogenic bacteria are more prevalent in gut microbiomes of diseased organisms, evidence from multiple animal models has shown that a lack of commensal bacteria can also contribute to the detrimental progression of various behavioral disorders (21, 22). In Autism Spectrum Disorder mouse models, the abundance of commensal bacteria species, *Lactobacillus reuteri* is significantly reduced in comparison to wild type mice (15). Further, *Lactobacillus plantarum* has been used as a probiotic in zebrafish models to reduce stress-related behaviors and prevent stress-induced microbiome dysbiosis (23). *L. plantarum* was discovered to modulate anxiety related behavior through the GABAergic and serotonergic pathways (23). Additionally, gut bacterial composition influences host behavior through the modulation of Brain-Derived Neurotrophic Factor (BDNF) levels and serotonin metabolism (24). For instance, probiotic treatment with *Lactobacillus rhamnosus* in zebrafish affects shoaling behavior and brain expression levels of *bdnf* and serotoninergic pathways both of which have been implicated in aggression and multiple psychiatric disorders (25, 26). In addition to influencing neurotransmitters levels, bacteria from the genera *Lactobacillus* can activate afferent neurons in the intestine that modulate pain sensation and actions of defensive behavior in response to stress (27). This evidence supports the notion that certain bacteria can influence the activity of neuromodulatory pathways responsible for social behavior. Collectively, these relationships between host gut bacteria and nervous system function support the notion that the gut microbiome regulates social activity by influencing neuromodulatory pathways responsible for social behavior.

Zebrafish social behavior has been studied extensively (28–32). When paired, zebrafish interact agonistically with conspecifics and quickly form stable dominance relationships consisting of dominant and submissive fish (28, 33). Once dominance forms, dominants have priority to food, shelter, and mates while submissive fish experience chronic stress (34). Although the neurobiological bases of zebrafish social aggression and their long-term impact on social activity have been investigated; the effects of social dominance on gut microbiome are less understood (35). This is of particular importance because the effects of chronic social stress on organisms are detrimental and have been adversely linked to anxiety-like behavior relevant for survival in many social species (2, 36–40). Specifically, reduction of glucocorticoid signaling activity (caused by insufficient hormone availability) in chronic stress models has been associated with stress-related disorders (41). Interestingly, the gut microbiota has recently been thought to modulate downstream glucocorticoid receptors in the hippocampus resulting in behavioral abnormalities highlighting the relationship between stress and the gut microbiome (42). The stability of zebrafish social dominance and ease of quantifying their aggressive social activity provides an opportunity to examine the brain-gut microbiome axes and the impact of social stress on the gut biome as dominance relationships are established.

The zebrafish gut microbiome has been examined in embryos and adults. A past study showed that the gut microbiome composition of host zebrafish changes throughout its development (43). Specifically, as embryonic zebrafish develop, the diversity of gut bacteria decreases significantly in terms of the number of operational taxonomic units (OTUs), which are clusters of bacteria that exhibit high sequence similarity of the 16S rRNA gene (44, 45). However, microbiome composition was similar between stages of adult fish, which indicates that major gut microbial changes occur before and during major developmental changes such as sexual differentiation (44). The adult zebrafish gut microbiome is composed primarily of bacteria within the Proteobacteria, Firmicutes, and Fusobacteria phyla (46). Pathogenic bacteria within the Proteobacteria phylum are found in significantly lower relative abundance of healthy human gut microbiomes and higher relative abundance of Proteobacterial members can be used as a diagnostic for disease and dysbiosis (47). Additionally, in zebrafish, Proteobacteria including pathogenic genera *Vibrio* and *Plesiomona* were reduced in probiotic treated fish as opposed to commensal Firmicutes bacterial members (24). Thus, the presence of pathogenic bacteria in both zebrafish and humans can indicate microbial dysbiosis and potential health issues. In this study, we examined whether social dominance has opposing effects on gut microbiome in dominant versus chronically stressed submissive animals. We tested the hypothesis that the relative abundance, diversity, and overall composition of zebrafish gut microbiome is socially regulated. We also sought to determine whether inherent differences in gut microbiome prior to dominance formation would predict future social rank. We compared gut microbiomes (based on amplicon sequencing of the 16S rRNA gene) among socially dominant, submissive, socially isolated, and group-housed communal fish. Results from this work improve our understanding of how social aggression and stress impact host gut microbiome and may help future studies to understand the gut-brain axis and how the onset of behavioral disorders affect gut dysbiosis.

### Experimental Methods

#### Zebrafish husbandry

Zebrafish used in this study were wild-type (WT) fish between the age of 8-12 months old. Male and female zebrafish were housed communally (20 fish per tank) in an automated flow-through tank system (Aquatic Habitats Z-Hab System) in (33 × 21 × 19 cm) holding tanks with room light cycle (14h light/10h dark). Fish were fed a consistent diet of brine shrimp and pellets twice daily. All methods and protocols were approved by East Carolina University’s Institutional Animal Care and Use Committee (IACUC) AUP# D320b.

#### Pairing and behavioral analysis

Male zebrafish siblings (n=24) were isolated in individual tanks for 7 days to minimize the effects of prior social experience. All fish used in the experiment were taken from a wild-type communal. After initial isolation, 12 fish were randomly paired and placed into new experimental tanks while 6 fish remained in isolation, and the remaining 6 fish were placed into one new tank to serve as communal controls. All experimental tanks were (26 × 14 × 10 cm). Over a period of 14 days, the 6 zebrafish pairs self-established rank and their behavior was monitored and recorded over a course of 14 days (2 weeks). Selected fish were similar in size and behavior was observed visually, while written observations were made regarding dominant and subordinate behavior among the paired and communal fish. Although communal fish do not establish structured dominance hierarchies, periodic aggressive activities are observed. Specifically, the number of attacks and retreats by each fish were recorded during a 5-minute observation period 3 times a week. Additionally, tank locations of each fish (top vs. bottom of tank) were monitored and recorded as stressed or subordinate fish tend to stay lower in the tank and more aggressive/dominant fish claim the top of the tank.

#### Fecal sample collection

Initial fecal samples were collected from each fish on day 3 of the isolation period. This fecal collection before pairing or establishment of behaviors served as a method of comparison for later fecal collections. After pairing of fish after the 7-day isolation period, fecal samples were collected on day 0 (day of pairing), day 7 (one week after pairing), and day 14 (two weeks after pairing). Collections were made throughout the progression of social hierarchal development to determine whether the gut microbiome composition of host fish evolves as dominance was established.

On days of fecal collection, fish were removed from tanks and each fish was placed into individual containers to prevent any cross contamination of fecal matter. Before pairing, each fish was carefully observed and distinct markings, size differences, and color differences were noted to ensure easy recognition of each fish upon isolation and fecal sample collection. Two to four hours after separation and feeding each fish, fecal samples were collected (48, 49). Fecal samples from each fish were extracted using sterile micropipette tips and were placed into 500 microliters of DNAse free water. Fecal samples were extracted from the container of each individual fish soon after defecation to minimize cross contamination with the water in the container. This method of fecal sample collection has been utilized in previous studies examining gut microbiome composition. This method of fecal collection was selected as opposed to intestinal tract extraction (50, 51) in order to analyze microbiome composition over several time periods. Gloves were always worn when handling fecal samples and fish, to minimize any cross contamination. After samples were collected, they were stored at -80 °C until DNA extractions and processing were performed.

#### Microbiome analysis

To determine differences in microbial composition in zebrafish based on social status, microbial communities in each fecal sample were characterized with Illumina sequencing of the highly conserved 16S rRNA gene (52). Following fecal sample collection, DNA extractions were performed on each sample using the PowerLyzer PowerSoil DNA protocol. Approximately 0.02g of feces were collected from each fish for extractions. This extracted genomic DNA was used as the template for PCR reactions, where barcoded primers (515F-806RB) were used to target the V4 region of the 16S rRNA gene (52, 53). Each fecal sample was run in triplicate PCR reactions and then combined and purified using the Axygen® AxyPrep magnetic bead purification kit (Corning Life Sciences). After successful cleanup, DNA concentrations (ng/mL) from each sample were quantified using the Quant-iT dsDNA HS (high sensitivity) assay (Thermo Scientific, Waltham, Massachusetts, USA). Recorded DNA concentrations (ng/mL) using HS measurements were converted to the final DNA concentration (ng/μl) and then dilutions using PCR grade water were performed to ensure all samples added to the final pooled product were the same mass (ng). PCR products were pooled in equimolar concentrations and sequenced on an Illumina MiSeq platform using paired-end reads (Illumina Reagent Kit v2, 500 reaction kit) at the Center for Genomics and Bioinformatics at Indiana University.

We processed raw 16S rRNA gene sequences using a standard mothur pipeline (v1.40.1) (54, 55). We assembled contigs from paired end reads and quality trimmed using a moving average quality score of 35 bp. We aligned sequences to the SILVA rRNA gene database (v.128) (56), and we removed chimeric sequences using the VSEARCH algorithm (57). We divided sequences based on taxonomic class and binned into operational taxonomic units (OTUs) with a 97% sequence similarity level, and we classified OTUs using the SILVA rRNA database (v128). We used an OTU-based 3% distance threshold to avoid splitting the same bacterial genome into distinct clusters (using amplicon sequence variant (ASV) of a single base difference). In addition, the sequencing was conducted using short-reads (250 bp) and not assembled genomes, which introduces PCR and sequencing errors which can result in over-inflated number of ASVs (58). The broadscale community composition patterns are robust when employing an OTU-based approach (59).

#### Species diversity and community compositional analysis

We examined the diversity and composition of bacterial communities in each fecal sample. For each microbiome sample, we calculated the Chao1 OTU richness using the estimateR() function and Shannon diversity using the diversity() function in the vegan package (60), and also calculated Simpson’s evenness using a custom function after we rarefied the OTU table to 10,181 observations. We calculated the Bray-Curtis dissimilarity matrix and visualized community composition according to ‘Social Status’ and ‘Day’ using principal Coordinate Analysis (PCoA). Finally, genus level compositions with relative abundances greater than 0.05 were plotted after OTU table was rarefied to 6,000 observations.

#### Statistical analyses

All data analyses were completed using the R statistical environment (R v4.2.0, R Core Development Team 2022; R Studio v2022.07.1). Using the lmer() function from the lmerTest package (61), a linear mixed effects model with ‘Day’ and ‘Social Status’ as fixed effects was used to analyze bacterial diversity metrics Chao1 OTU richness, Shannon diversity (H’), and Simpson’s evenness. We ran a permutational multivariate analysis of variance (PERMANOVA) based on the Bray-Curtis dissimilarity of bacterial community composition to determine the extent that ‘Social Status’ and ‘Day’ and the interaction explained bacterial community composition. We next identified bacterial species that represented each treatment (‘Social Status x ‘Day’) for bacterial taxa with a relative abundance greater than 0.05 when summed across all samples. We performed the PERMANOVA using the adonis() function from the vegan package (60) and the indval() function in the indicspecies package (62).

#### Data Availability

All code and data used in this study are in a public GitHub repository (https://github.com/PeraltaLab/ZebrafishMicrobiomes_SocialStatus) and NCBI SRA BioProject PRJNA925886.

## RESULTS

### Bacterial species diversity and community composition

We compared gut microbiomes of zebrafish of different social statuses. Bacterial OTU richness (Fig. 1) was similar across ‘Social Status’ and ‘Day’, while the interaction of ‘Social Status’ and ‘Day’ but not the main effects influenced Shannon diversity (Fig. 2) and Simpson’s evenness increased from Day 7 to Day 14 (Fig. 3) (Table S1). While statistical significance was detected, the linear models explained a relatively low amount of variation (richness: adjusted R^2^= 0.149, p-value=0.020; diversity: adjusted R^2^=0.106, p-value=0.062; evenness: adjusted R^2^=0.215, p-value=0.003) (Table S1). Communal data points plotted similarly to isolate data points during the isolation period (IP) before any pairing or rank establishment. Additionally, dominants data points trended similarly to those of subordinate animals during the isolation period. These trends were observed for richness (Fig. 1) and evenness (Fig. 3).

**Figure 1.**
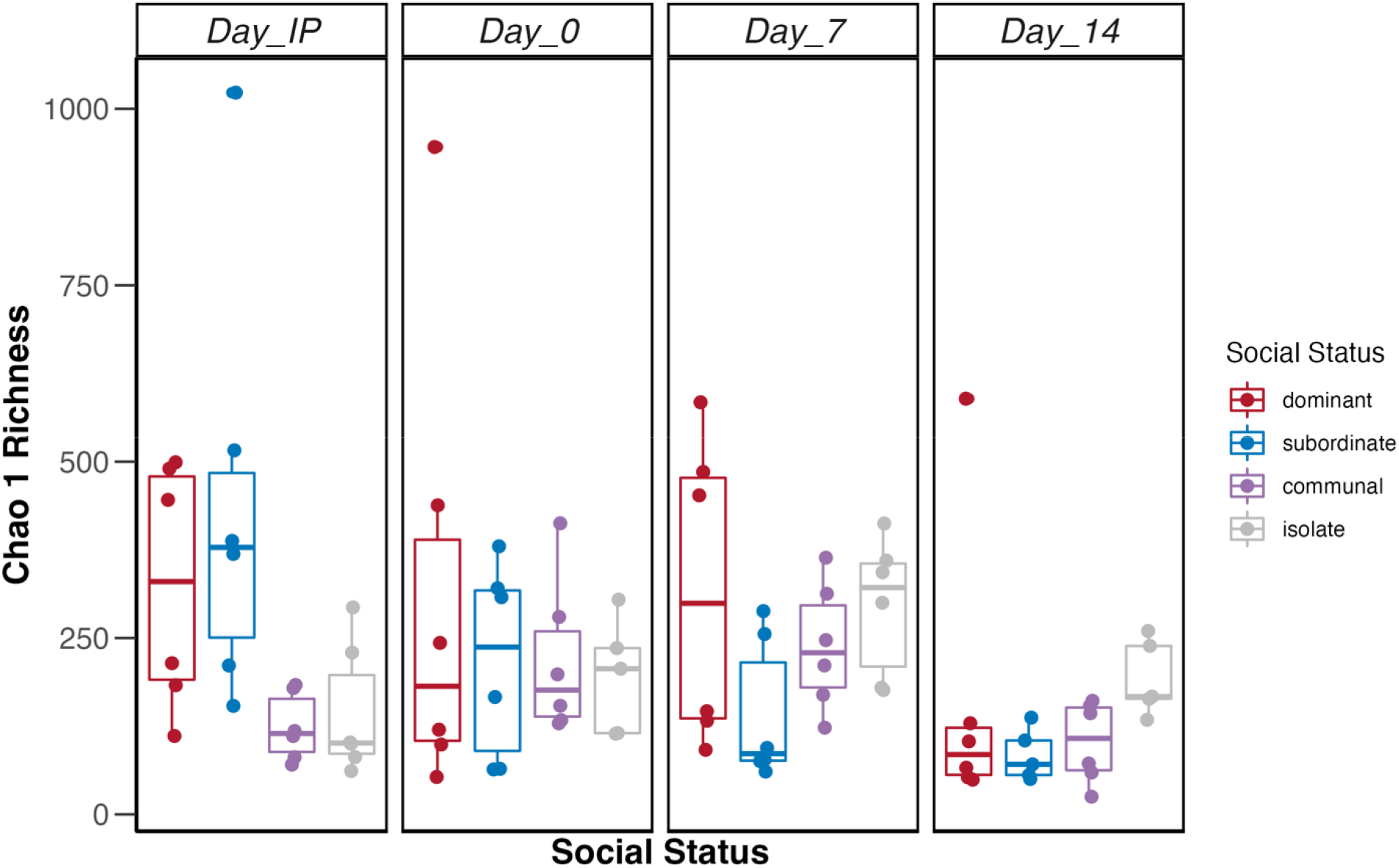
Boxplots representing bacterial Chao 1 richness in individuals from dominant (red), subordinate (blue), communal (purple), and isolate (gray) animals throughout the isolation period (Day IP) representing collection before pairing and pairing period (Day 0, Day 7, Day 14). The boxplot is a visual representation of 5 key summary statistics: the median, the 25% and 75% percentiles, and the whiskers which represent the feasible range of the data as determined by 1.5 × the interquartile range. Symbols represent individual raw data points from six replicate samples. Summary of statistical output in Appendix S1: Table S1A.

**Figure 2.**
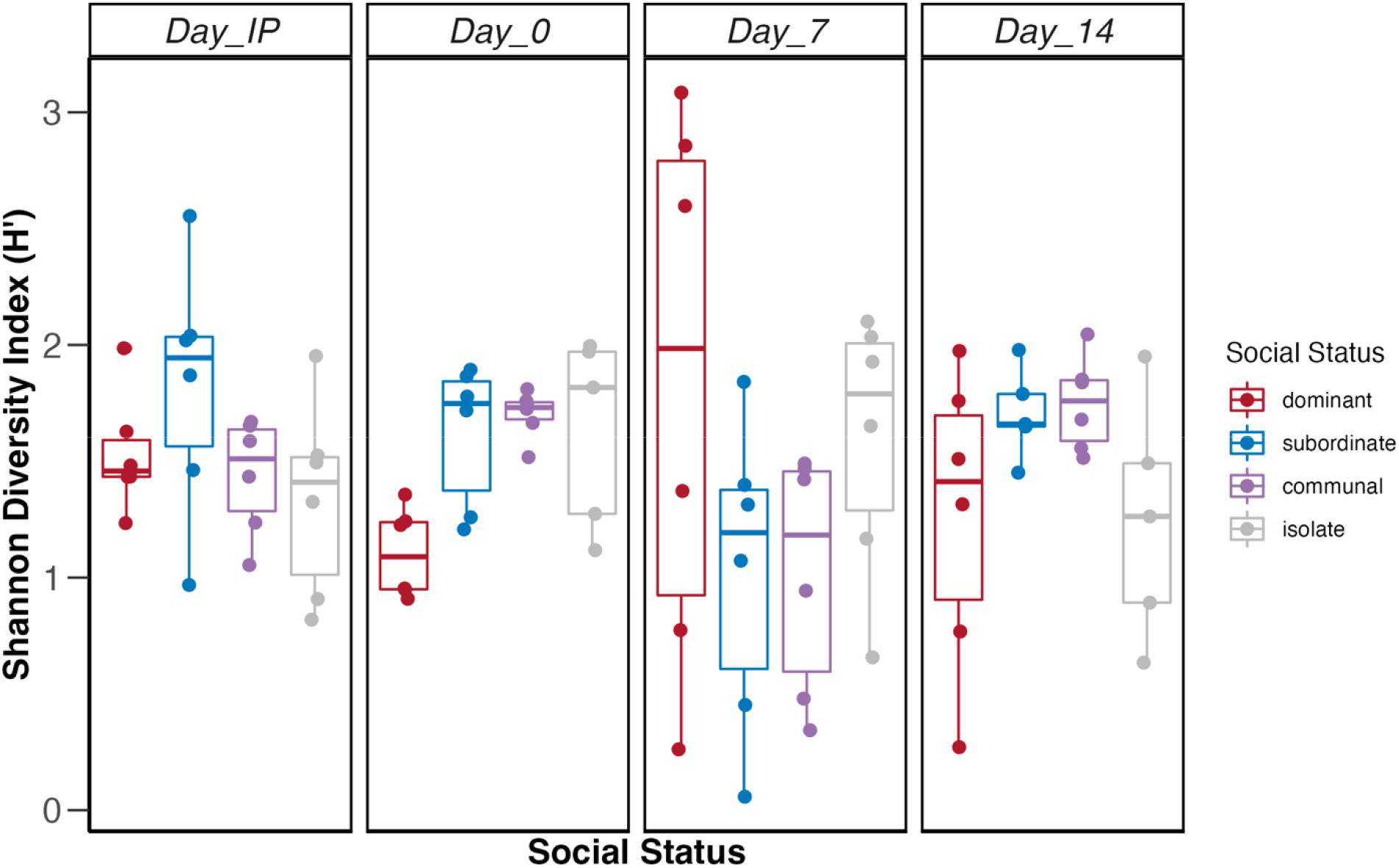
Boxplots representing bacterial Shannon diversity index in individuals from dominant (red), subordinate (blue), communal (purple), and isolate (gray) animals throughout the isolation period (Day IP) representing collection before pairing and pairing period (Day 0, Day 7, Day 14). The boxplot is a visual representation of 5 key summary statistics: the median, the 25% and 75% percentiles, and the whiskers which represent the feasible range of the data as determined by 1.5 × the interquartile range. Symbols represent individual raw data points from six replicate samples. Summary of statistical output in Appendix S1: Table S1B.

**Figure 3.**
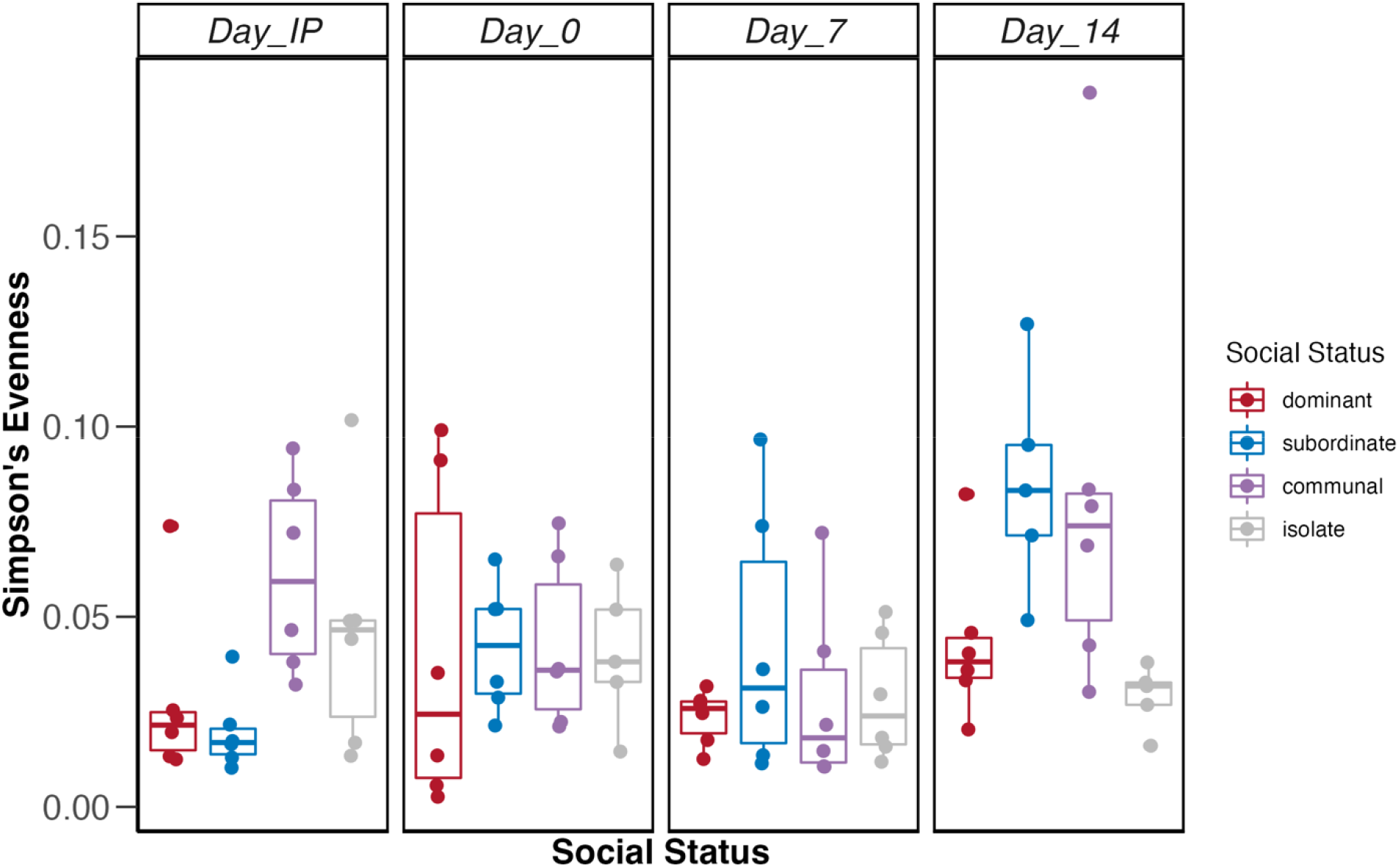
Boxplots representing bacterial Simpson’s diversity index in individuals from dominant (red), subordinate (blue), communal (purple), and isolate (gray) animals throughout the isolation period (Day IP) representing collection before pairing and pairing period (Day 0, Day 7, Day 14). The boxplot is a visual representation of 5 key summary statistics: the median, the 25% and 75% percentiles, and the whiskers which represent the feasible range of the data as determined by 1.5 × the interquartile range. Symbols represent individual raw data points from six replicate samples. Summary of statistical output in Appendix S1: Table S1B.

In contrast to bacterial diversity, ‘Social Status’ and ‘Day’ influenced bacterial community composition. The interaction of ‘Social Status’ and ‘Day’ accounted for 14.4% of the variation in bacterial composition (F_9,92_=1.9214, P=0.001), while the main effects of ‘Day’ accounted for 12.5% (F_3,92_=5.035, P=0.001) and ‘Social Status’ accounted for 9.1% (F_3,92_=3.671, P=0.001) of the bacterial community composition (Fig. 4, Table S2). The bacterial gut microbiomes of dominant and isolate fish tended to group together while the gut microbiomes of communal fish were distinct from all other groups, except for Day 14 which grouped by the gut microbiomes of subordinate fish on Day 0 (Fig. 4). The isolation period (IP) for subordinate, dominant, and isolated fish resulted in similar gut microbiomes (diamond symbols). For subordinate and dominant fish (blue and red symbols), the gut microbiomes were more variable over time (i.e., symbols were farther apart over ordination space) more than the isolate and communal gut microbiomes (gray and purple symbols) (Fig. 4).

**Figure 4.**
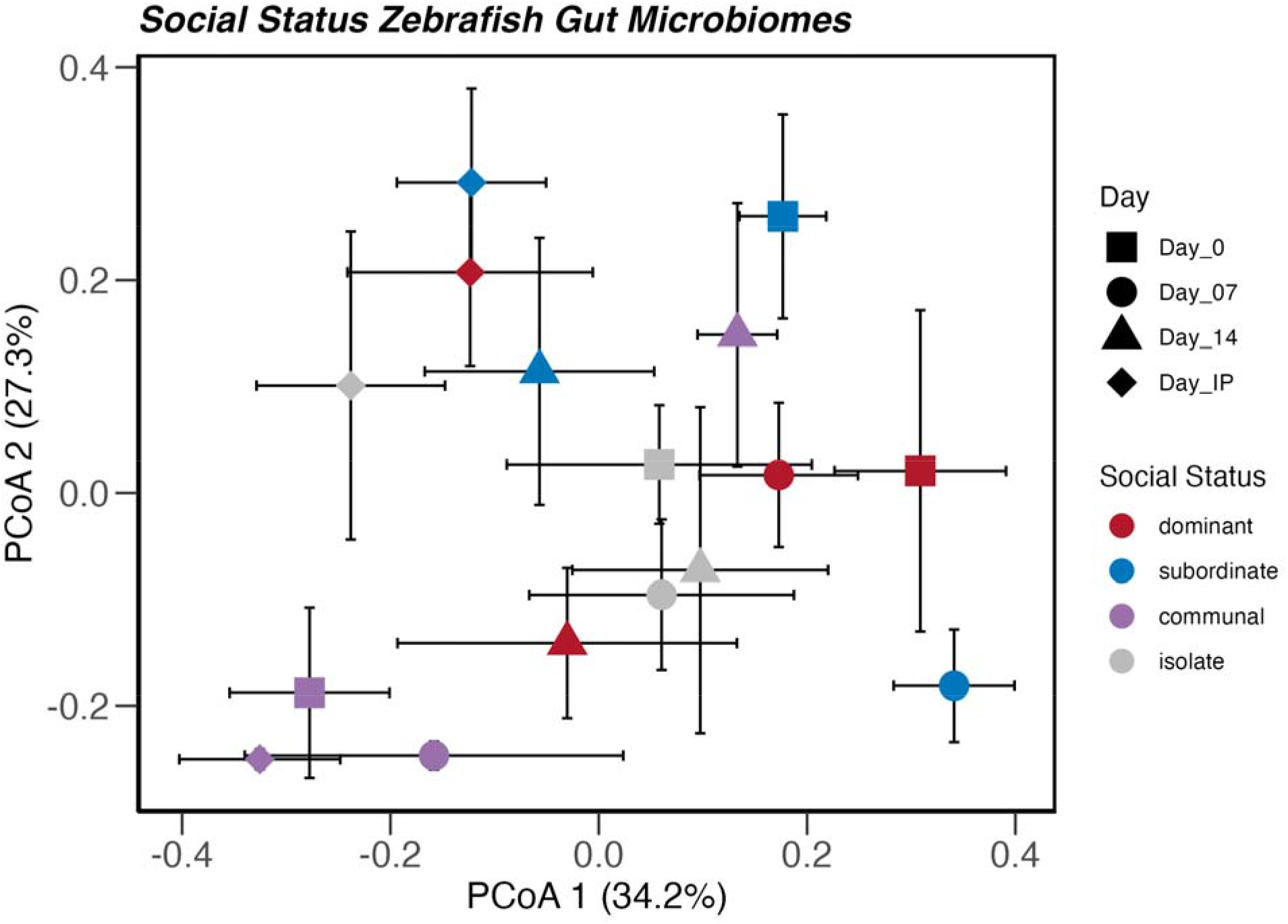
Ordination based on a Principal Coordinates Analysis depicting bacterial community composition according to social status and day. Symbols are colored according to social status (dominant=red, subordinate=blue, communal=purple, and isolate=gray) and shapes represent day of pairing (Day 0=square, Day 7=circle, Day 14=triangle, Day IP=diamond). The centroid and standard error bars (along axes 1 and 2) were calculated for six replicate plots. Summary of statistical output in Appendix S1: Table S2.

### Taxa-level Shifts Across Social Status

We evaluated which bacteria taxa represented gut microbiomes by ‘Social Status’ and ‘Day’ using indicator species analysis. *Paracoccus spp*. and *Achromobacter spp*. represented gut microbiomes of dominant fish during the isolation period, and unclassified taxa from the families Planctomycetacea, Caldilineaceae, Comamonadaceae and indicators *Staphylococcus spp*. and *Exiguobacterium spp*. represented gut microbiomes of dominant fish on Day 7 (Table S3). *Acinetobacter spp*. and *Psychrobacter spp*. represented gut microbiomes of subordinate fish during Day 0 and Day 7, respectively (Table S3). Unclassified taxa from the class Betaproteobacteria and family Rhizobiaceae represented gut microbiomes of isolated fish on Day 0 (Table S3). For communal fish, an unclassified taxon from the Rhodobacteraceae family, *Pseudomonas spp*., and *Stappia spp*. represented gut microbiomes during the isolation period, while *Rhizobiales spp*., *Arthrobacter spp*., and *Acinetobacter spp*. represented Day 0, *Chryseobacterium spp*. represented Day 7, and *Shewanella spp*. represented Day 14 (Table S3). Unclassified taxa from the class Betaproteobacteria and family Rhizobiaceae represented gut microbiomes of isolated fish on Day 0 (Table S3).

To further explore trends, we evaluated genus level composition for genera observed in > 5% relative abundance. While community composition varied between different social statuses and day of pairing, we observed changes in dominant individuals on day 7 and 14 of pairing (Fig. S1). We observed a reduction in pathogenic genera that were consistently high in relative abundance in non-dominant social status groups and throughout the pairing period. Specifically, *Aeromonas* and unclassified Enterobacteriaceae relative abundance was lower in dominant individuals on day 7 and 14, whereas relative abundance of these genera remained higher in other groups (Fig. S1). We also observed an increased relative abundance of *Chitinobacteria, Vibrio*, and *Pseudomonas spp*. in subordinate, isolate, and communal animals and the presence of these genera were maintained before and during pairing (Fig. S1). In addition, certain commensal genera such as *Exiguobacterium* and *Cetobacterium* increased in relative abundance in dominant animals on day 7 of pairing. These genera were present in dominant animals throughout pairing, but on day 7, the relative abundance increased, especially for the *Cetobacterium* genus (Fig. S1).

## Discussion

A growing number of studies highlight the microbial role in the link between the gut and the brain (63–65). Our results suggest a link between the host gut microbiome and social experience in zebrafish building on prior research that gut microbiome may be associated with social activity and behavioral disorders. Here, we studied whether social aggression and chronic stress influence gut microbiome content in zebrafish. We showed that while there were no significant differences in gut microbiome diversity or richness among different social states, social dominance leads to compositional differences. This result complements prior work in a mouse model that showed a causative relationship between the gut and anxiety/depression phenotypes (15). These similarities along with findings in other studies examining how probiotic treatment alters behavioral activity in socially stressed individuals lends further credence of the strong relationship between the gut-brain axis. Specifically, chronically stressed zebrafish that were treated with probiotics, experienced a complete reversal and rescue of their anxiety-like behavior (23). This supports a microbiome mediated behavior hypothesis in which changes to the gut microbiome affects associated behaviors related to stress and anxiety. However, in those studies, both the composition and diversity of the gut microbiome directly affects behavior whereas we found that only bacterial composition was affected by social status.

While we observed compositional differences between fish of different social rank, these differences were more pronounced after 7-14 days of social interactions when social dominance solidified. This signifies that the chronic stress that accompanies social submission or isolation may directly impact bacterial composition. Additionally, it seems that social dominance and aggression promote an increased presence of commensal bacterial genera. Previous research in autism models show an inverse relationship between the gut and behavior, as individuals born with gut microbiota dysbiosis are predisposed to social deficits and autism-like behaviors (66).

One of the questions we sought to answer in this study was whether inherent differences in gut microbiome prior to social dominance formation would predispose individuals to become either dominant or submissive. Based on our results of fecal samples that were collected prior to pairing during the isolation period (Day IP), this hypothesis is not supported in this model. However, it would be interesting to determine in future studies whether repeated probiotic or antibiotic treatments over extended periods are sufficient to modulate aggression levels and reverse social dominance. Such results would be very exciting and would strongly indicate that the microbiome could directly affect social aggression.

Based on previous research in zebrafish and other animal models, the genus, *Lactobacillus*, from phylum Bacteroidetes, is a common protective bacterial species often found in the guts of many animal models (23). Because of these protective qualities, the *Lactobacillus* genus is often used as a probiotic treatment in both zebrafish and mouse models to rescue stressed phenotypes (15, 23). Interestingly, the *Lactobacillus* genus was not present in the taxonomic analysis of this experiment. This is likely due to *Lactobacillus* species being a transient to the gut microbiome, as they primarily colonize the host GI tract from food intake and the oral cavity (67–69). The only genus of bacteria found from the Bacteroidetes phylum was *Chryseobacterium spp. a*nd was primarily found in communal fish on day 7 (Table S3). While this genus doesn’t come from the same family as *Lactobacillus*, both genera seem to have similar protective qualities in zebrafish gut microbiota. In a study examining bacterial key players in pathogenic infection in zebrafish, *Chryseobacterium massillae* was determined to be important in protecting both larvae and adult zebrafish from pathogenic infection (70). Interestingly, this bacterial genus was only present in low relative abundance in both communal and dominant individuals (Table S3, Fig. S1).

Within the Firmicutes phylum, the bacterial genus *Exiguobacterium* was also determined as a component of compositional analysis of the zebrafish gut in our study. This genus of bacteria has been shown to produce cyclic dipeptides, which have many antimicrobial, antifungal, antiviral, and anti-inflammatory properties in humans and other animals (71, 72). Intriguingly, *Exiguobacterium acetylicum* has therapeutic properties in zebrafish colorectal cancer models, which highlights the protective properties of this genus of bacteria in the gut of both zebrafish and humans (73). While the bacterial genus *Exiguobacterium* was present in socially submissive and isolated zebrafish, this OTU only increased in relative abundance in dominant individuals after a week of pairing (Fig. S1). The absence of the bacterial genus *Exiguobacterium* in communal animals should be noted, which may be due to the heterogeneity among mid-ranked members living in a communal setting whereby individuals may dominate some low-ranking animals while being submissive to high-ranking opponents.

Given the high percent of Proteobacteria communities in zebrafish gut microbiomes, the presence of pathogenic bacteria is not surprising, particularly since a large portion of the species in these potential pathogenic genera are waterborne. Therefore, tank water was analyzed and indicated the tank water microbiome tested in this study grouped in the middle of subordinate, dominant, isolate, and communal fish microbiomes (Figure S2). However, the varying relative abundance of certain potential pathogenic bacteria like *Aeromonas*, unclassified Enterobacteriaceae, *Staphylococcus, Vibrio*, and *Pseudomonas spp*. in the community composition is noteworthy. *Aeromonas and* unclassified Enterobacteriaceae were the dominant putatively pathogenic genera in all groups and at all time points during pairing, but several putatively pathogenic genera were primarily seen in subordinate, isolate, and even communal individuals (Fig. S1). Many of these genera have been used in zebrafish infection models. A model of *Staphylococcus* infection in zebrafish has been studied, and the interactions between *Staphylococcus aureus* and other bacteria in the gut to create infection (74). This study specifically examined the immune response of the fish in response to this infection, but it is important to note the pathogenesis associated with *Staphylococcus* infection. Interestingly, *Staphylococcus* relative abundance is increased in dominant individuals even though other pathogenic genera are decreased. This could contribute to the increase in Firmicutes relative abundance seen in dominant individuals. This finding is inconsistent with other trends observed, as other potential pathogenic bacterial communities decreased in dominant individuals; however, these observations still provide insight into gut microbiome composition and associated social experience. In addition to the presence of *Staphylococcus* in dominant individuals, indicator species analysis also revealed the presence of *Achromobacter* and *Paracoccus* in dominant individuals during the isolation period (Table S3). *Achromobacter* and *Paracoccus* are genera of bacteria with pathogenic properties in both mammals and fish (75– 77). While these genera are present in dominant individuals prior to pairing, they decrease in relative abundance post-pairing which supports the hypothesis that social status may affect gut bacterial composition. Prior studies did not describe the role for the unclassified genera indicators observed for dominant day 7, which included OTUs in *Planctomycetaceae, Caldineaceae*, and *Comamondaceae* genera. The *Aeromonas* genus, which has a high relative abundance in almost all zebrafish individuals regardless of social status, has been observed in several pathogenic models of zebrafish. Interestingly, specific species within the *Aeromonas* genus have been seen to induce severe infection of zebrafish models by influencing the release of reactive oxygen species (ROS) and nitrogen free radicals triggering an immune response accompanied by massive inflammatory reaction and increased mortality rates (78). While our results do not specify the species of *Aeromonas* present, higher relative abundances of this genus may result in pathogenesis in chronically stressed animals as well (e.g., socially submissive or isolates) (79, 80). We also acknowledge the limitations to 16S rRNA gene sequencing and discuss the compositional differences with emphasis on putative pathogenic and commensal genera. While we observe notable shifts in both commensal and pathogenic bacteria abundance between fish of different social conditions across time, we acknowledge the effect on host health depends on the interactions within microbial community and interactions between host and its microbiomes. To further identify how these specific genera may be involved in the development of social status, future studies examining the changes to specific commensal and pathogenic genera and their effects on social status are needed. This could be accomplished through experimental antibiotic additions or to specific manipulation of bacterial taxa via probiotic additions. In mice, antibiotic treatment has resulted in depressive phenotypes and gut microbiome dysbiosis (81). On the other hand, probiotic treatments have shown a rescue of anxiety-like and stressed behaviors in both mice and zebrafish (23, 24, 82). The addition of probiotic *Lactobacillus casei* increased survival of zebrafish larvae infected with *Aeromonas veronii* (83). Altogether, these previous research findings support the effectiveness of both antibiotic addition and probiotic manipulation in examining social conditions, behavioral disorders, and development.

Collectively, our results show that gut microbiome composition is socially regulated. This is illustrated by a partial shift in the composition from pathogenic to commensal bacteria in dominant animals; while chronically stressed animals (e.g., submissive, and socially isolated) showed an increase in potentially pathogenic bacteria and a decline in commensal bacterial genera. The interconnectivity between gut function and cognitive processes are likely to mediated bi-directionally via afferent and efferent neural and neuroendocrine signaling pathways (82). Although our results suggest that the physiological mechanisms underlying the shift in microbial composition is slow acting and require one to two weeks for their biological manifestation, prior reports have shown that acute exposure to stress can impact the gut microbiota community composition by changing the relative proportions of the main microbiota phyla (85). Moreover, experimental manipulations of gut microbiota affect stress levels and anxiety-like behavior by influencing the function of the hypothalamic-pituitary-adrenal stress axis (86). Indeed, social stress induced increase in blood cortisol levels correlates with changes in the diversity of the mammalian gut microbiome, including the abundance of lactic acid and pathogenic bacteria (85, 87). In addition, chronic injections of glucocorticoids can have positive and negative impact on the presence of specific microbial taxa, which consequently affect host metabolism (88–90). Thus, it is likely that chronic stress induced by either prolonged social isolation or subordination likely to have increased cortisol levels in isolated and submissive zebrafish with substantial increase in expression of gut pathogenic bacteria. Chronic stress has also been shown to negatively impact the immune response in zebrafish and other model organisms (91–93). It would be beneficial to examine the effects of social dominance on the immune response in zebrafish to determine if the gut microbiome composition and/or social conditions play a role in altered immune functions.

How these status-dependent changes in gut bacterial composition feedback to influence brain function remains poorly understood. Given that zebrafish experiencing induced stress exhibit changes in gut microbiome composition and brain gene expression (24), future experiments testing this relationship in zebrafish in the context of social dominance and stress will be highly informative. More specifically, it would be of interest to determine whether transcriptional changes in the brains of dominant and subordinate fish can be linked to changes in bacterial abundance in the host gut. Indeed, a large body of work has shown that dopaminergic, GABAergic, and serotoninergic pathways implicated in social behavior are highly plastic, but whether the plasticity is mediated by feedback regulation of gut microbiome to influence motivational brain centers involved in social aggression remains an open question (32, 94–98).

## Supporting information

Supplemental_Materials_Zebrafish_Microbiomes_ScottE

## ACKNOWLEDGMENTS

We thank Dr. Katie Clements for her assistance in the early design of behavioral experiments. This work was supported by the National Science Foundation grant (#1754513) to F.A.I. and grant (#1845845) to A.L.P..

