## Supplemental_Materials_Zebrafish_Microbiomes_ScottE for "The effects of social experience on host gut microbiome in male zebrafish (*Danio rerio*)"

Table S1. Analysis of variance table comparing bacterial Shannon diversity index (A) and Simpson’s diversity index (B) due to social status (dominant, submissive, communal, isolate) and day (Day 0, Day 7, Day 14, Day IP).

(A) Bacterial Chao 1 richness

|  | DF | SumSq | MeanSq | F-value | P-value |
| --- | --- | --- | --- | --- | --- |
| Social.Status | 3 | 149770 | 49923 | 1.854 | 0.145 |
| Day | 3 | 219518 | 73173 | 2.717 | 0.050 |
| Social.Status:Day | 9 | 467909 | 51990 | 1.930 | 0.060 |
| Residuals | 77 | 2073843 | 26933 |  |  |

Adjusted R-squared: 0.149, F-statistic: 2.072 on 15 and 77 DF, p-value: 0.020

(B) Bacterial Shannon diversity index

|  | DF | SumSq | MeanSq | F-value | P-value |
| --- | --- | --- | --- | --- | --- |
| Social.Status | 3 | 0.137 | 0.046 | 0.176 | 0.913 |
| Day | 3 | 0.404 | 0.134 | 0.519 | 0.671 |
| Social.Status:Day | 9 | 6.187 | 0.687 | 2.651 | 0.010 |
| Residuals | 77 | 19.965 | 0.259 |  |  |

Adjusted R-squared: 0.106, F-statistic: 1.730 on 15 and 77 DF, p-value: 0.062

(C) Bacterial Simpson’s diversity index

|  | DF | SumSq | MeanSq | F-value | P-value |
| --- | --- | --- | --- | --- | --- |
| Social.Status | 3 | 0.0065 | 0.002 | 2.573 | 0.060 |
| Day | 3 | 0.010 | 0.003 | 4.530 | 0.006 |
| Social.Status:Day | 9 | 0.014 | 0.002 | 2.091 | 0.040 |
| Residuals | 77 | 0.058 | 0.001 |  |  |

Adjusted R-squared: 0.215, F-statistic: 2.675 on 15 and 77 DF, p-value: 0.003

Table S2. Summary PERMANOVA comparing bacterial communities due to social status (dominant, submissive, communal, isolate) and day (Day 0, Day 7, Day 14, DayIP).

| Factor | DF | SumOfSqs | R^2^ | F-value | P-value |
| --- | --- | --- | --- | --- | --- |
| Social.Status | 3 | 2.301 | 0.091 | 3.671 | 0.001 |
| Day | 3 | 3.156 | 0.125 | 5.035 | 0.001 |
| Social.Status:Day | 9 | 3.614 | 0.144 | 1.921 | 0.001 |
| Residual | 77 | 16.092 | 0.639 |  |  |
| Total | 92 | 25.163 | 1 |  |  |

Table S3. Bacterial taxa (OTUs) representing the unique taxa associated to treatment type (Social Stats x Day) according to indicator species analysis.

| OTU | Cluster | IndVal | Prob | Phylum/Class/Order/Family/Genus |
| --- | --- | --- | --- | --- |
| Otu00057 | Dominant.Day_7 | 0.712 | 0.002 | Planctomycetes/Planctomycetia/Planctomycetales/Planctomycetaceae/Planctomycetaceae_unclassified |
| Otu00063 | Dominant.Day_7 | 0.540 | 0.002 | Chloroflexi/Caldilineae/Caldilineales/Caldilineaceae/Caldilineaceae_unclassified |
| Otu00047 | Dominant.Day_7 | 0.536 | 0.023 | Proteobacteria/Betaproteobacteria/Burkholderiales/Comamonadaceae/Comamonadaceae_unclassified |
| Otu00013 | Dominant.Day_7 | 0.440 | 0.015 | Firmicutes/Bacilli/Bacillales/Staphylococcaceae/Staphylococcus |
| Otu00012 | Dominant.Day_7 | 0.357 | 0.048 | Firmicutes/Bacilli/Bacillales/Bacillales_Incertae_Sedis_XII/Exiguobacterium |
| Otu00028 | Dominant.Day_IP | 0.512 | 0.010 | Proteobacteria/Alphaproteobacteria/Rhodobacterales/Rhodobacteraceae/Paracoccus |
| Otu00037 | Dominant.Day_IP | 0.219 | 0.045 | Proteobacteria/Betaproteobacteria/Burkholderiales/Alcaligenaceae/Achromobacter |
| Otu00017 | Subordinate.Day_0 | 0.483 | 0.023 | Proteobacteria/Gammaproteobacteria/Pseudomonadales/Moraxellaceae/Acinetobacter |
| Otu00045 | Subordinate.Day_7 | 0.430 | 0.005 | Proteobacteria/Gammaproteobacteria/Pseudomonadales/Moraxellaceae/Psychrobacter |
| Otu00021 | Communal.Day_0 | 0.336 | 0.006 | Proteobacteria/Alphaproteobacteria/Rhizobiales/Rhizobiales_unclassified/Rhizobiales_unclassified |
| Otu00027 | Communal.Day_0 | 0.317 | 0.002 | Actinobacteria/Actinobacteria/Actinomycetales/Micrococcaceae/Arthrobacter |
| Otu00009 | Communal.Day_0 | 0.251 | 0.024 | Proteobacteria/Gammaproteobacteria/Pseudomonadales/Moraxellaceae/Acinetobacter |
| Otu00011 | Communal.Day_14 | 0.196 | 0.048 | Proteobacteria/Gammaproteobacteria/Alteromonadales/Shewanellaceae/Shewanella |
| Otu00019 | Communal.Day_7 | 0.520 | 0.008 | Bacteroidetes/Flavobacteriia/Flavobacteriales/Flavobacteriaceae/Chryseobacterium |
| Otu00030 | Communal.Day_IP | 0.498 | 0.002 | Proteobacteria/Alphaproteobacteria/Rhodobacterales/Rhodobacteraceae/Rhodobacteraceae_unclassified |
| Otu00004 | Communal.Day_IP | 0.299 | 0.002 | Proteobacteria/Gammaproteobacteria/Pseudomonadales/Pseudomonadaceae/Pseudomonas |
| Otu00024 | Communal.Day_IP | 0.270 | 0.011 | Proteobacteria/Alphaproteobacteria/Rhodobacterales/Rhodobacteraceae/Stappia |
| Otu00020 | Isolate.Day_0 | 0.592 | 0.001 | Proteobacteria/Betaproteobacteria/Betaproteobacteria_unclassified/Betaproteobacteria_unclassified/Betaproteobacteria_unclassified |
| Otu00022 | Isolate.Day_0 | 0.388 | 0.016 | Proteobacteria/Alphaproteobacteria/Rhizobiales/Rhizobiaceae/Rhizobiaceae_unclassified |

Figure S1. Relative abundance for bacterial genera representing > 5% of the gut microbiome across Social Status (communal, dominant, subordinate, isolate) and sampling period (isolation period (IP), Day=0, Day=7, Day=14). Colors represent different genera.

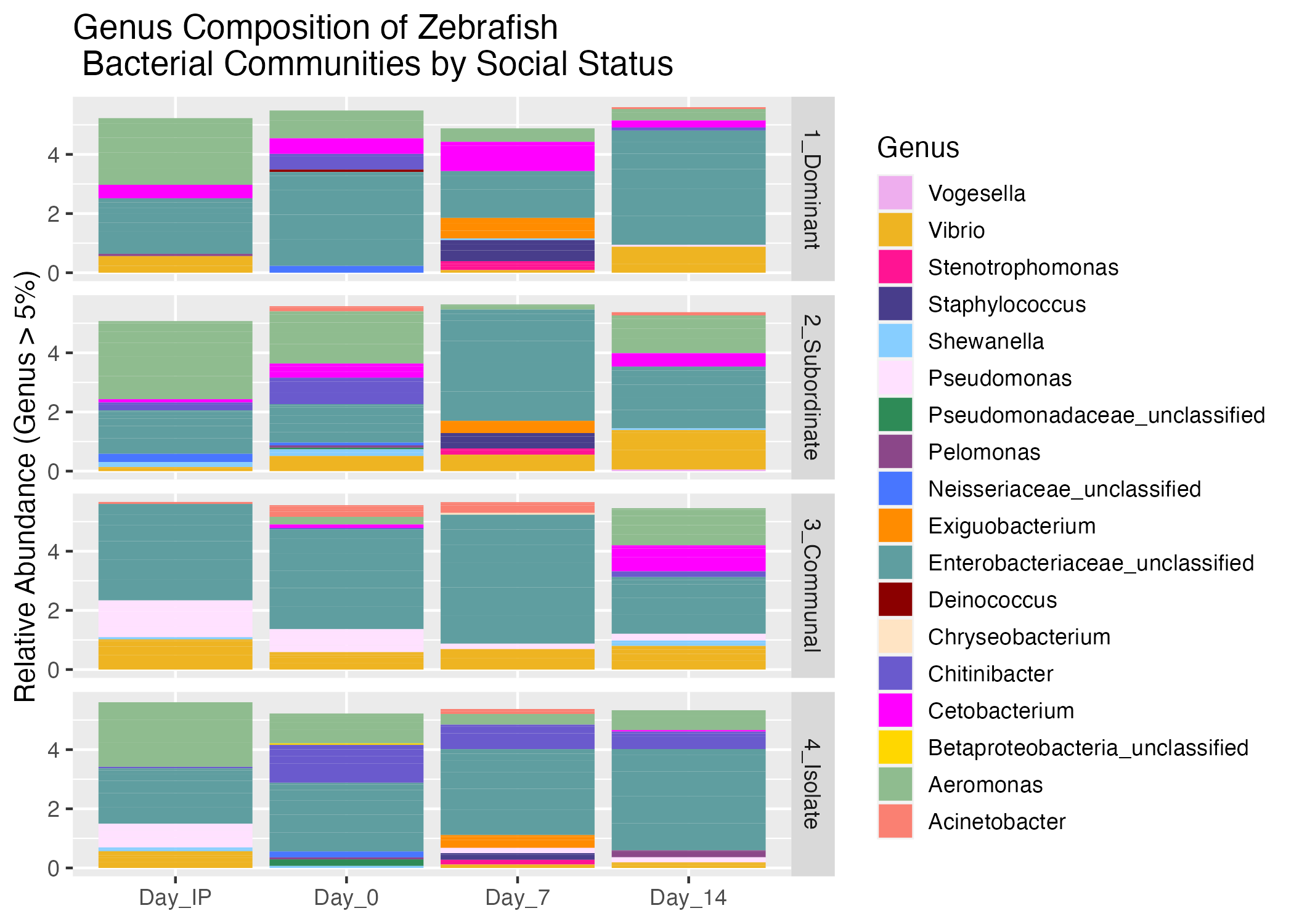

Figure S2. Ordination based on a Principal Coordinates Analysis depicting bacterial community composition according to social status and day. Symbols are colored according to social status (dominant=red, subordinate=blue, communal=purple, isolate=gray, water=black), and shapes represent day of pairing (Day 0=square, Day 7=circle, Day 1=triangle, Day 14=diamond, Day IP=circle). The centroid and standard error bars (along axes 1 and 2) were calculated for six replicate plots for fish microbiomes but only a single replicate was represented for the tank water environmental microbiome.

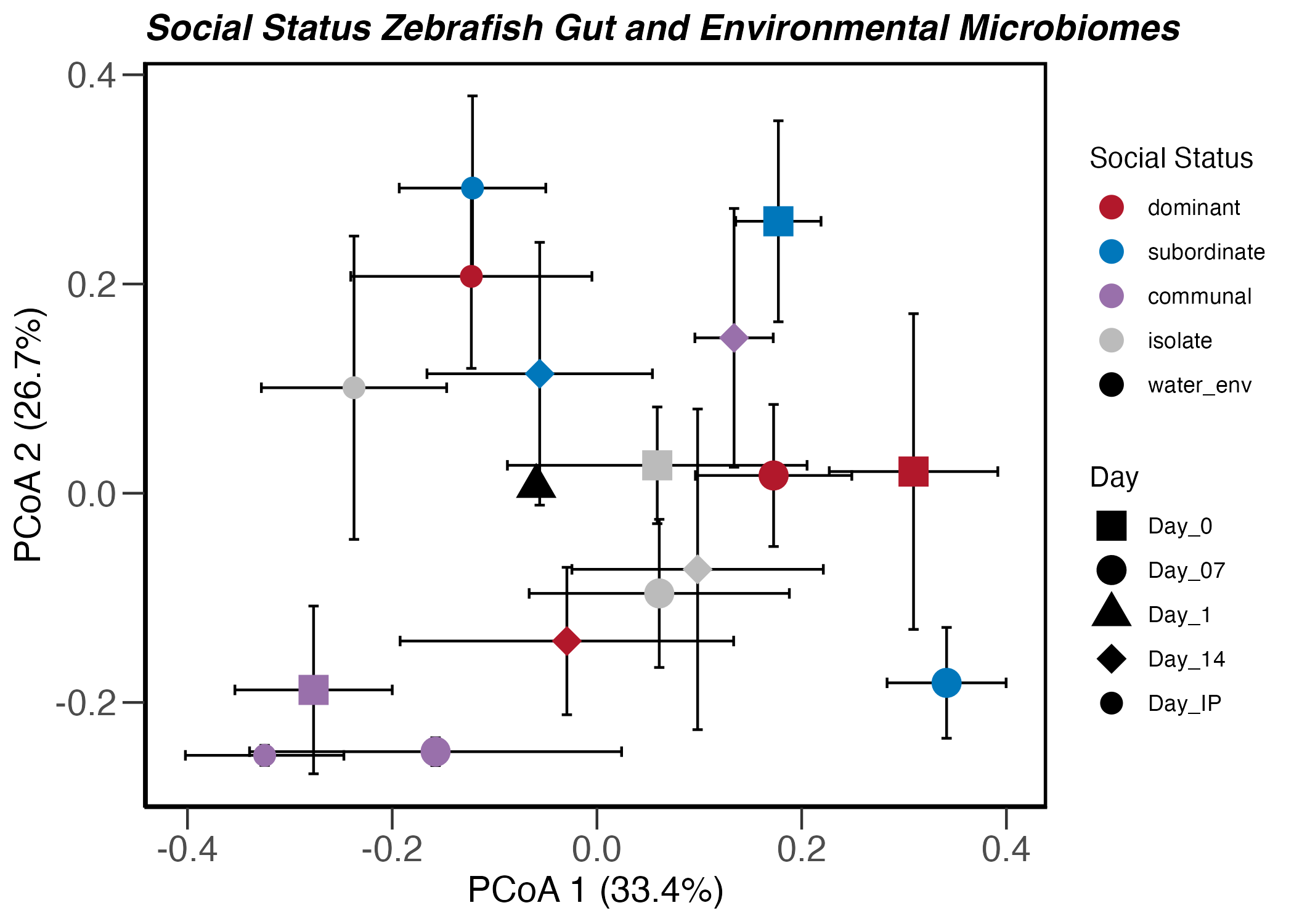
